# Social dysfunction relates to altered default mode network functional integrity across neuropsychiatric disorders: A replication and generalization study

**DOI:** 10.1101/2025.01.09.631642

**Authors:** Simon Braak, Maarten Mennes, Tanja Su, Yolande Pijnenburg, Geor Bakker, Celso Arango, Nic van der Wee, Stéphanie Bauduin, Ana Ortiz-Tallo Moya, Javier-David Lopez-Morinigo, Gerard R. Dawson, Abigail B. Abrahams, Amy Christine Beckenstrom, Christian F. Beckmann, Hugh M. Marston, Brenda W.J.H. Penninx, Martien J. H. Kas

## Abstract

**Background:** Social dysfunction is an early manifestation of neuropsychiatric disorders that may relate to altered Default Mode Network (DMN) integrity. This study aimed to replicate previous findings linking social dysfunction with diminished resting-state DMN functional connectivity and altered task-based DMN functional activation in response to emotional faces across schizophrenia (SZ), Alzheimer’s disease (AD), and healthy controls (HC), and to extend these findings to major depressive disorder (MDD).

**Methods:** Resting-state fMRI and task-based fMRI data on implicit facial emotional processing were acquired in an overlapping cohort (resting-state fMRI: N=167; SZ=32, MDD=44, AD=29, HC=62. Task-based fMRI: N=152; SZ=30, MDD=42, AD=26, HC=54). Additionally, mega-analyses (N=317 for resting-state fMRI; N=291 for task-based fMRI) of the current and a prior independent sample were conducted. Social dysfunction was indexed with the Social Functioning Scale (SFS) and the De Jong-Gierveld Loneliness (LON) scale.

**Results:** The association between higher mean SFS+LON social dysfunction scores and diminished DMN connectivity within the dorsomedial prefrontal cortex across SZ/AD/HC participants was replicated, and extended to MDD patients. Similar observations within the dorsomedial and rostromedial prefrontal cortex were found in the mega-analysis. Associations between social dysfunction and DMN activation in response to sad and happy faces were not replicated or found in the mega-analysis.

**Conclusions:** Diminished dorsomedial prefrontal cortex DMN connectivity emerged as a transdiagnostic neurobiological marker for social dysfunction, suggesting a potential treatment target for precision medicine approaches. DMN functional responses to emotional faces may not be a sensitive biomarker for social dysfunction.

## Introduction

Social dysfunction is an early common manifestation of several neuropsychiatric disorders, including schizophrenia (SZ), major depressive disorder (MDD), and Alzheimer’s disease (AD), that strongly affects patient health outcomes (1–3). Despite presenting with distinct core symptoms, patients across these disorders may have similar social dysfunction and abnormalities in brain networks that govern social behaviors (3–6). In other words, social dysfunction may behave as a transdiagnostic feature across neuropsychiatric disorders. Understanding the shared neurobiological factors associated with social dysfunction across these disorders may lead to novel targeted treatments. This aligns with the Research Domain Criteria framework, which aims to investigate mental disorders within the context of major domains of neuro-behavioral functioning, including the social processes domain, relevant for multiple disorders (7). In this regard, the entire continuum, from normal social functioning to severe dysfunction, can be studied both in healthy controls (HC) and in patients across disorders (7), which has been indicated as ‘transdiagnostic’ in previous studies (6, 8, 9).

Emerging findings support the presence of transdiagnostic neural correlates of social dysfunction, particularly alterations in Default Mode Network (DMN) connectional integrity (8, 10–13). Specifically, the Psychiatric Ratings using Intermediate Stratified Markers 1 (PRISM1) study demonstrated that behavioral aspects of social dysfunction (e.g., social withdrawal and interpersonal dysfunction) and perceived loneliness were transdiagnostically associated with diminished intrinsic functional connectivity within the rostromedial prefrontal cortex (rmPFC) and dorsomedial PFC (dmPFC) across SZ and AD patients and HC (8). In line with this, a composite score of behavioral and subjective aspects of social dysfunction (i.e., small social network, perceived social disability, perceived loneliness) has been linked to diminished intrinsic functional connectivity within the rmPFC and posterior superior frontal gyrus in MDD patients (10). Similar observations in autism spectrum disorder, Attention Deficit/Hyperactivity Disorder (ADHD), and 22q11.2 deletion syndrome, suggest that this neural correlate of social dysfunction may be independent of diagnosis (14–16).

Additionally, aberrant functional activation of the DMN in response to emotional face stimuli has been observed across various populations and is linked to social dysfunction (12, 13, 17, 18). The PRISM1 study found that, across 46 SZ and 40 AD patients and 53 HC, behavioral aspects of social dysfunction were associated with greater functional activation of key nodes of the DMN (e.g., dmPFC and angular gyrus) in response to sad faces (12). This was accompanied with reduced functional activation in a subset of these brain regions (e.g., angular gyrus) in response to happy faces (12). Consistent with this, greater functional activation of the left angular gyrus in response to sad faces was linked to perceived social disability across 200 persons with and without depressive and/or anxiety disorders, suggesting that social dysfunction might converge predominantly on sad emotional face processing of the DMN across neuropsychiatric disorders (13). Although such associations were not observed for perceived loneliness in both studies, other research has linked neural processing of emotional faces to perceived loneliness, albeit not in a transdiagnostic manner (12, 13, 19, 20).

Nevertheless, it remains to be investigated whether the neurobiological factors associated with social dysfunction reported by the PRISM1 study are reproducible in an independent sample and extend to other neuropsychiatric disorders. This is crucial for developing targeted treatment strategies aimed at the shared neuro-pathophysiology underlying social dysfunction, independent of diagnostic classifications. Therefore, in the current study, we examined the reproducibility of the association between social dysfunction and diminished resting-state DMN functional connectivity and altered functional activation of the DMN in response to sad and happy faces across SZ/AD patients and HC, as well as whether these associations may extend to MDD patients. We hypothesize that greater severity of behavioral aspects of social dysfunction and perceived loneliness would be associated with diminished DMN functional connectivity of the medial PFC across SZ/AD patients and HC (8). Furthermore, we hypothesize that, across SZ/AD/HC participants, greater severity of behavioral aspects of social dysfunction will be associated with greater functional activation within DMN regions in response to sad faces, along with reduced activation in these brain regions in response to happy faces (12). Finally, it was anticipated that our findings would extend to MDD patients and be independent of diagnosis (10, 13).

## Methods and Materials

### Participants

Data were acquired in the PRISM2 study, which investigates the replicability and generalizability of the PRISM1 study (21) in a new independent sample. Participants were recruited between June 2022 and December 2023 from two sites in the Netherlands (Amsterdam UMC and Leiden University Medical Centre), and two sites in Spain (Hospital General Universitario Gregorio Marañón & Hospital Universitario De La Princesa). Data collection followed harmonized protocols across sites, in line with the PRISM1 study (8, 12, 22). The study was approved by the local Medical Ethical Review Boards and participants provided written informed consent.

Inclusion/exclusion criteria aligned with the PRISM1 study (8, 12). SZ participants (age: 18– 45) required DSM-IV diagnosis, <15 years since diagnosis, and positive and negative syndrome scale (PANSS) positive symptoms score <22 (23). MDD participants (age: 18–55) required DSM-IV diagnosis without psychotic features and a Quick Inventory of Depressive Symptomatology - Self-Rated (QIDS-SR) score > 5 (24). AD diagnosis (age: 50–80) followed NIA-AA criteria with Mini Mental State Examination scores (MMSE-2) of 20–26 (25, 26). All patients (SZ, MDD, AD) had to be stabilized on their central nervous medication for at least eight weeks. Two HC groups were recruited: HC-younger (age: 18–55) and HC-older (age: 56–80), with exclusions for any Axis-I psychiatric or cognitive-impairing neurological diagnoses. Further details are in the Supplement.

### Social dysfunction assessment

Social dysfunction was assessed using two self-report questionnaires: the Social Functioning Scale (SFS) and the De Jong-Gierveld Loneliness (LON) scale (27, 28), in line with PRISM1 (8, 12). The SFS measures behavioral aspects of social dysfunction and consists of seven subscales: social withdrawal, interpersonal functioning, prosocial activities, recreational activities, independence-competence, independence-performance, and employment (27). Employment items were excluded due to high rates of retirement in the AD and HC-older group. The SFS scores were reversed so higher scores indicated greater social dysfunction, with the total score used in the imaging analyses. The LON scale was used to index the subjective experiences of social dysfunction (i.e., perceived loneliness) (28). Responses were classified according to the scoring guidelines to indicate the presence or absence of loneliness symptoms, yielding a total score ranging from 0 to 11, where higher scores indicated greater perceived loneliness. The SFS and LON scales are psychometrically validated to reliably measure behavioural aspects of social functioning (27, 29) and perceived loneliness (28, 30) across neuropsychiatric disorders. The two scales show a moderate to strong correlation (Spearman’s r = 0.69, *p*<0.0001), indicating they assess partly distinct aspects of social dysfunction.

### Imaging paradigms

Consistent with PRISM1, resting-state data were acquired by instructing participants to lie still and look at the fixation cross displayed in the middle of the screen without falling asleep (8). The implicit facial emotional processing (iFEP) fMRI task (Figure 1) was used to investigate functional activation in response to sad, happy, and fearful emotional face stimuli (see Supplement for details) (12).

**Figure 1.**
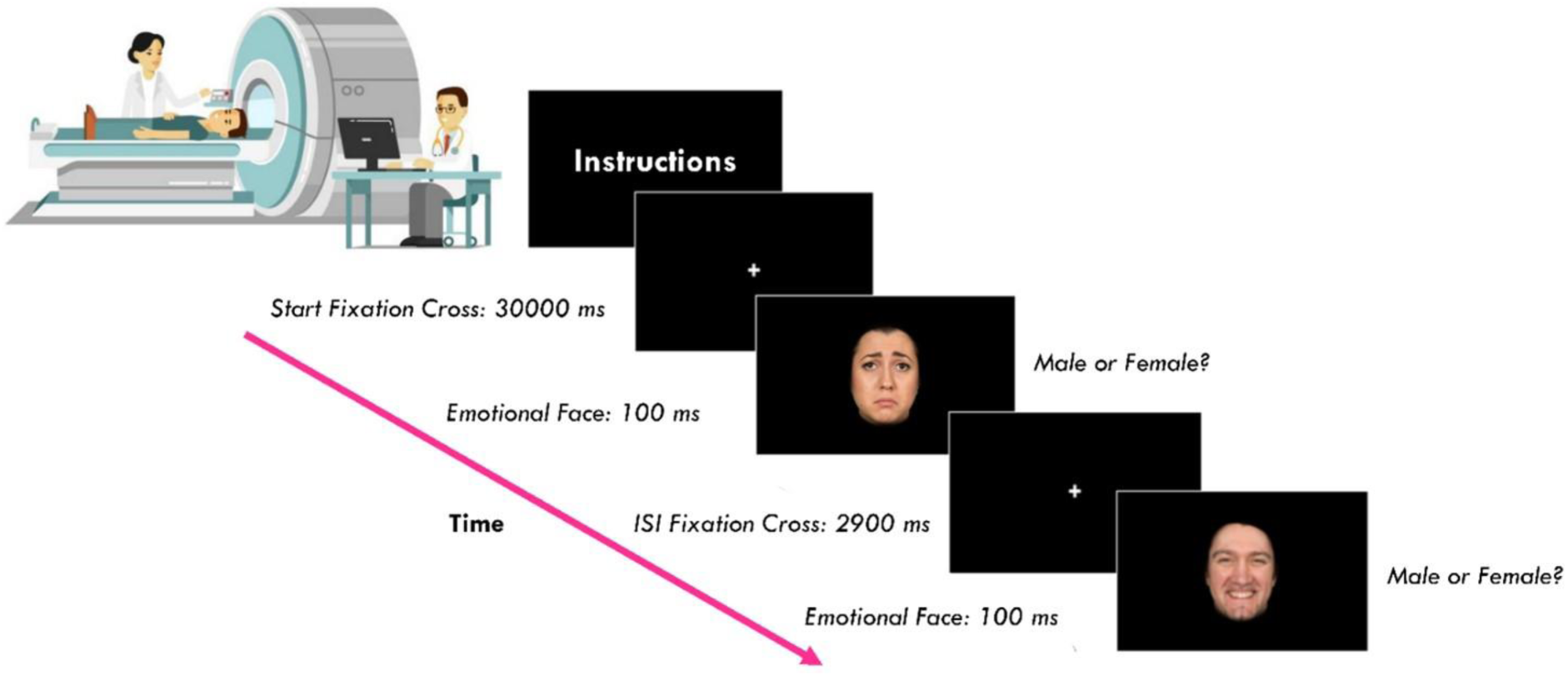
Time course of stimulus presentation for the implicit facial emotional processing task during the scanning session. ISI = interstimulus interval. Figure adapted from Braak et al., 2024 (12). Emotional faces are reproduced with permission from P1vital Products Ltd.

### MRI data acquisition and preprocessing

Imaging data were acquired using harmonized protocols and the same parameters as PRISM1 (see Supplement for acquisition details) (8, 12). A Philips Achieva 3T MRI scanner (Leiden University Medical Centre) and Philips Ingenia 3T MRI scanner (Amsterdam UMC) with a 32-channel head coil were used at the Dutch sites. MRI assessments for both Spanish recruiting sites were performed on a single Siemens Prisma 3T MRI scanner with a 64-channel head coil. Imaging data were preprocessed using FMRIB Software Library (FSL) version 6.0.7.6 (31), following established protocols used in PRISM1 (see Supplement) (8, 12).

### Statistical Analyses

Demographic and clinical characteristics for the five groups (SZ, MDD, AD, HC-younger, HC-older) were compared in R (version 4.2.1) using χ2 for dichotomous variables and analysis of variance for continuous variables with Tukeýs method as a post-hoc test. If parametric test assumptions were not met the Kruskal-Wallis test was applied, followed by Dunn’s test for post-hoc analysis.

#### Resting-state functional connectivity analyses

The analysis pipeline for resting-state data followed previously established methods in PRISM1, focussing on whole-network functional connectivity within the DMN (see Supplement) (Figure 2) (8). Using FSL’s MELODIC module, two group-level independent component analysis (ICA) DMN maps were generated: one for SZ/AD/HC participants and another including the MDD participants. Spatial similarity of the DMN maps was assessed using Pearson’s r.

**Figure 2.**
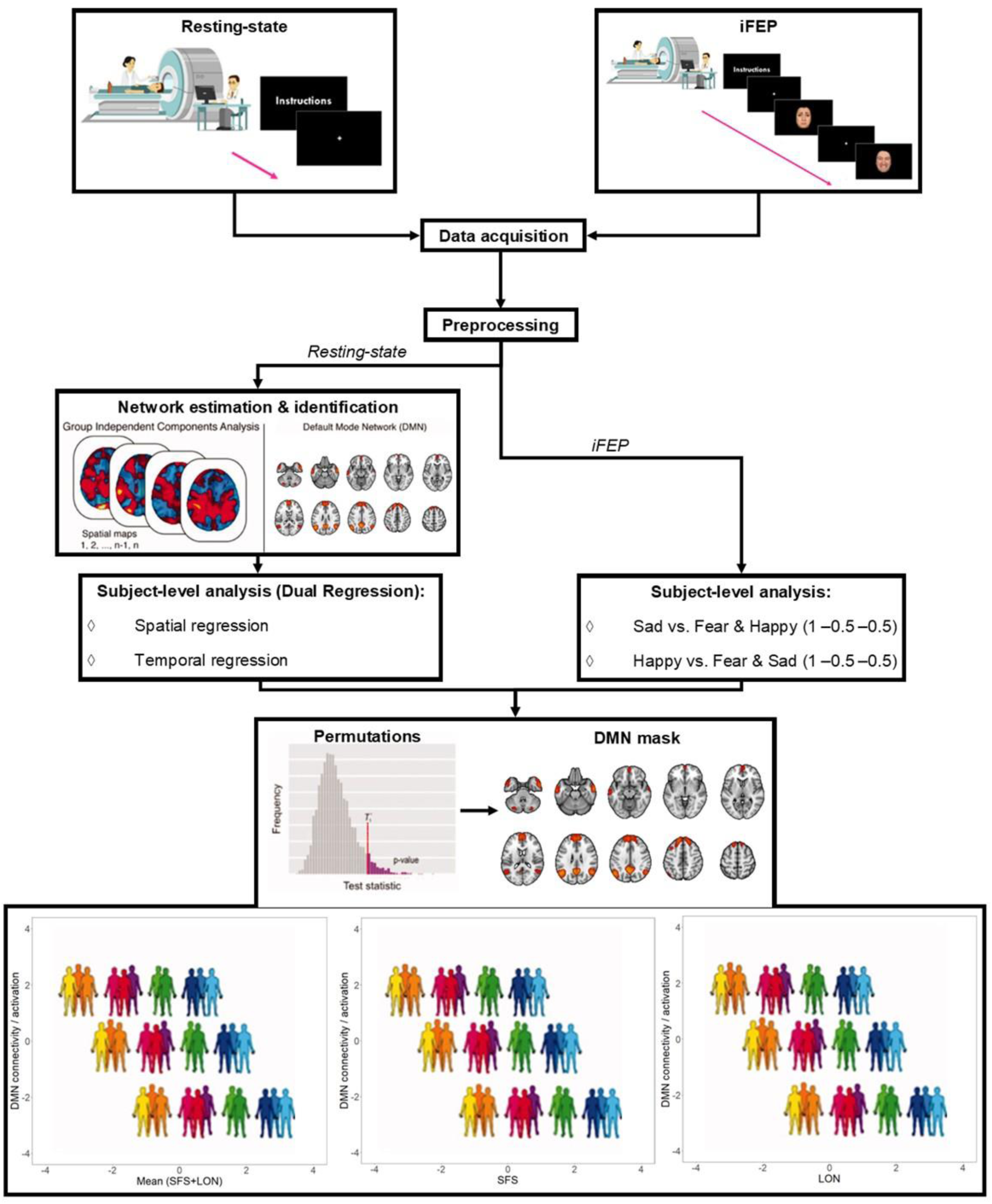
Functional connectivity and activation analyses of the Default Mode Network. Resting-state fMRI data and task-based implicit facial emotional processing (iFEP) fMRI data was collected and preprocessed. Using FSL’s MELODIC module, two group-level independent component analyses (ICA) were performed: one for schizophrenia, Alzheimer’s disease, and healthy control participants, and another including the major depressive disorder participants. These sets of spatial maps from the group-average ICA were used to generate subject-specific versions of the spatial maps, and associated timeseries, using dual regression. The Default Mode Network (DMN) maps were selected based on its topological architecture. Regarding iFEP data, two contrasts were tested in the subject-level General Linear Models to measure neural responses to sad and happy emotional faces. Finally, associations between DMN intrinsic functional connectivity or functional activation in response to sad and happy emotional faces and the social dysfunction indicators were analyzed using FSL’s Randomise tool with N=5000 permutations. As the DMN was the primary focus of the current study, the statistical output of the iFEP data was pre-threshold masked with the DMN map of the group-average resting-state ICA obtained with the total sample including MDD participants. For completeness, these analyses were also run at a whole-brain level. Statistical thresholding and correction for multiple comparisons achieved through Threshold-Free Cluster Enhancement with family-wise error correction at *p*<0.05. Adapted with permission from Wiley Periodicals, Inc.: Human Brain Mapping (32). Emotional faces are reproduced with permission from P1vital Products Ltd.

Associations between diminished DMN functional connectivity and higher SFS and/or LON total scores were analyzed using general linear model (GLM) analyses with FSL’s Randomise tool, controlling for clinical (diagnostic status, psychotropic medication, comorbid symptomology) and sociodemographic factors (age, age squared, gender, education, scan type). The GLM included the SFS and LON scores as separate regressors, wherein the cumulative, as well as the unique, effect of these constructs on DMN connectional integrity was examined. These analyses were applied to the samples with and without the MDD group. Post-hoc network-specificity analyses were run to examine whether brain-social behavior associations could be found within the salience network and central executive network (CEN) (see Supplement). This approach aligns with the triple network model of psychopathology, which posits that disrupted functional integrity within and between the DMN, salience network, and CEN increases susceptibility to various clinical manifestations, including social dysfunction (33). Finally, a mega-analysis of pooled PRISM1 and PRISM2 data was conducted to confirm effects, enhance robustness, and reduce Type I error by increasing statistical power and precision (8). A pooled group-level ICA DMN connectivity map was generated for all the data from PRISM 1 and PRISM2 (including MDD participants) and associations with social dysfunction were examined in the same manner as the PRISM1 and PRISM2 studies, while additionally correcting for study sample (PRISM1/PRISM2). Imaging analyses were performed with all independent variables demeaned across participants, with statistical thresholding and correction for multiple comparisons achieved through Threshold-Free Cluster Enhancement (TFCE) with family-wise error (FWE) correction at *p<*0.05 (34).

#### Implicit facial emotional processing task analyses

The same iFEP data analysis pipeline was used as in prior work (12) and was performed on both the sample without and with MDD (see Supplement) (Figure 2). In short, two contrasts were tested in the subject-level GLM’s: Sad vs. Fear & Happy (1 -0.5 -0.5) and Happy vs. Fear & Sad (1 -0.5 -0.5). Group-level GLM analyses were performed using FSL’s Randomise tool on the subject-level statistical maps, correcting for the same covariates as in the resting-state analyses, and examining the cumulative and unique effects of SFS and LON scores on task-related neural activity (35, 36). As the DMN was the primary focus of the current study, the statistical output was pre-threshold masked with the DMN map of the group-average resting-state ICA obtained with the total sample including MDD. For completeness, analyses were also run at a whole-brain level. The mega-analysis of pooled PRISM1 and PRISM2 data was conducted in the same manner as the PRISM1 (12) and PRISM2 studies, additionally correcting for study sample (PRISM1/PRISM2). Statistical thresholding and correction for multiple comparisons was achieved through TFCE with FWE correction at *p<*0.05 (34).

## Results

### Patient characteristics

Participants were scanned for both resting-state fMRI and iFEP fMRI, with N=167 participants (SZ=32, MDD=44, AD=29, HC-younger=34, HC-older=28) for final resting-state analyses and N=152 participants (SZ=30, MDD=42, AD=26, HC-younger=29, HC-older=25) for final iFEP analyses. Characteristics of the resting-state and iFEP PRISM2 sample are provided in Table 1 and Table S1, respectively. Additionally, the mega-analysis samples are shown in Table S2 and Table S3. A flowchart detailing the number of participants for each analysis conducted is shown in Figure 3.

**Figure 3.**
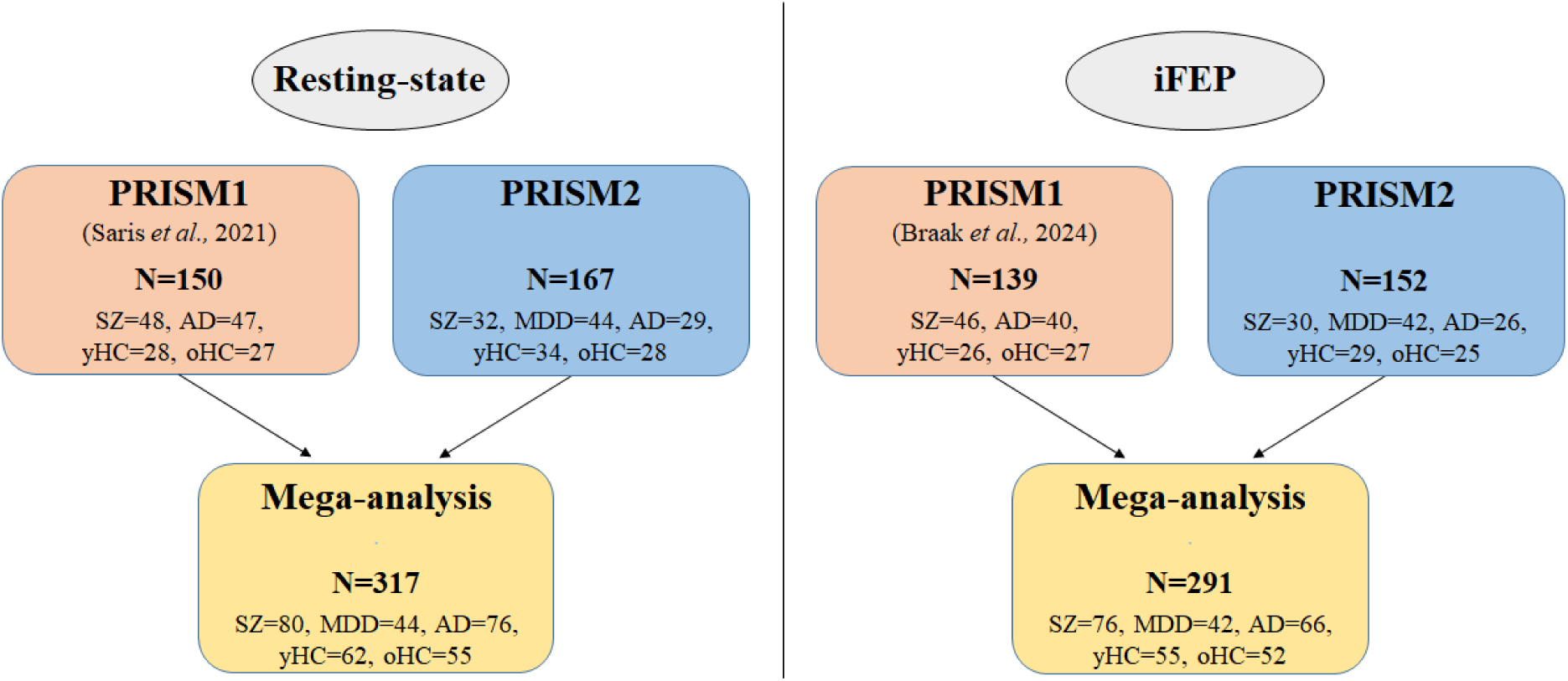
Flowchart detailing the number of participants for the imaging analyses. All participants in the PRISM1 and PRISM2 studies were scanned for both resting-state fMRI and task-based implicit facial emotional processing (iFEP) fMRI. SZ: schizophrenia, AD: Alzheimer’s disease. MDD: major depressive disorder, yHC: young healthy controls, oHC: older healthy controls.

**Table 1.** Resting-state sample characteristics of each group. Mean and standard deviations (SD) are displayed for continuous variables. When assumptions were violated, median and Q1 – Q3 are also displayed. Chi-square tests were performed for categorical variables. For continuous variables, analysis of variance with Tukeýs method as a post-hoc test was performed. In case assumptions were violated, a Kruskal-Wallis test with Dunńs test as a post-hoc test was performed. PANSS = Positive and Negative Syndrome Scale. MMSE = Mini-mental state examination. QIDS-SR = Quick Inventory of Depressive Symptomatology, Self-rated. SFS = Social Functioning Scale. A significance level of *p*<0.05 was considered statistically significant.

|  | Total Sample | Schizophrenia (SZ) | Major depressive disorder (MDD) | Alzheimer's disease (AD) | Young healthy controls (yHC) | Old healthy controls (oHC) | Pairwise differences |
| --- | --- | --- | --- | --- | --- | --- | --- |
|  | N=167 | N=32 | N=44 | N=29 | N=34 | N=28 |  |
| **Demographics** |  |  |  |  |  |  |  |
| Age (years), median (Q1 – Q3) | 44.0 (32.0 – 67.0) | 33.5 (30.0 – 38.3) | 39.0 (31.0 – 48.3) | 72.0 (68.0 – 77.0) | 33.5 (28.0 – 44.0) | 70.0 (65.3 – 74.0) | AD & oHC > SZ & MDD & yHC |
| *mean age (SD)* | 47.8 (18.6) | 33.3 (6.8) | 38.4 (11.7) | 71.6 (5.7) | 36.0 (11.1) | 69.1 (5.3) |  |
| Gender (% female) | 55.7% | 43.8% | 61.4% | 55.2% | 55.9% | 60.7% | $p > 0.05$ |
| Education (years), median (Q1 – Q3) | 15.0 (12.0 – 18.0) | 12.0 (10.0 – 16.0) | 14.5 (12.0 – 16.3) | 14.0 (12.0 – 20.0) | 17.0 (14.3 – 18.0) | 15.0 (12.8 – 18.0) | SZ < AD & yHC & oHC; MDD < yHC |
| *mean education (SD)* | 14.9 (3.8) | 12.9 (3.6) | 14.5 (3.7) | 15.3 (4.1) | 16.5 (3.0) | 15.5 (3.7) |  |
| Number of individuals per scan site |  |  |  |  |  |  |  |
| Amsterdam UMC | 53 | 11 | 14 | 11 | 8 | 9 |  |
| Leiden UMC | 32 | 5 | 11 | 0 | 11 | 5 |  |
| Spanish sites | 82 | 16 | 19 | 18 | 15 | 14 |  |
| Specific disorder characteristics |  |  |  |  |  |  |  |
| Psychotropic medication |  |  |  |  |  |  |  |
| Antipsychotic (%) | 17.4% | 90.6% | 0.0% | 0.0% | 0.0% | 0.0% |  |
| Antidepressant (%) | 23.4% | 12.5% | 65.9% | 20.7% | 0.0% | 0.0% |  |
| Acetylcholinesterase inhibitor and/or<br>NMDA receptor antagonist (%) | 10.8% | 0.0% | 0.0% | 62.1% | 0.0% | 0.0% |  |
| Benzodiazepines (%) | 9.6% | 15.6% | 15.9% | 13.8% | 0.0% | 0.0% |  |
| Antiepileptic (%) | 1.8% | 6.3% | 2.3% | 0.0% | 0.0% | 0.0% |  |
| Other psychotropics (%) | 6.0% | 6.3% | 15.9% | 3.4% | 0.0% | 0.0% |  |
| Severity disorder |  |  |  |  |  |  |  |
| Positive symptoms, PANSS, mean (SD) | NA | 11.5 (2.5) | NA | NA | NA | NA |  |
| Negative symptoms, PANSS, mean (SD) | NA | 14.9 (5.8) | NA | NA | NA | NA |  |
| MMSE, median (Q1 – Q3) | NA | NA | NA | 25.0 (23.0 – 25.0) | 29.5 (29.0 – 30.0) | 29.5 (29.0 – 30.0) | AD > yHC & oHC |
| *mean MMSE (SD)* |  |  |  | 23.6 (2.0) | 29.3 (0.8) | 29.2 (1.0) |  |
| QIDS-SR, median (Q1 – Q3) | 3.0 (2.0 – 10.0) | 5.0 (2.8 – 11.0) | 14.0 (10.0 – 19.0) | 2.0 (1.0 – 3.0) | 2.0 (0.3 – 2.0) | 2.0 (2.0 – 3.0) | MDD > SZ > AD & yHC & oHC |
| *mean QIDS-SR (SD)* | 6.2 (6.1) | 6.6 (4.7) | 13.9 (5.4) | 2.8 (2.1) | 1.9 (1.6) | 2.1 (1.1) |  |
| Social dysfunction scores |  |  |  |  |  |  |  |
| Reversed SFS score, median (Q1 – Q3) | 20.9 (14.8 – 29.9) | 24.4 (16.5 – 31.7) | 30.2 (25.3 – 36.6) | 25.3 (19.1 – 29.8) | 15.0 (9.3 – 17.6) | 13.6 (10.6 – 20.3) | MDD > AD & SZ > yHC & oHC |
| *mean SFS score (SD)* | 22.7 (10.6) | 24.9 (10.2) | 31.2 (8.7) | 24.3 (9.3) | 14.0 (5.3) | 15.5 (7.1) |  |
| Loneliness score, median (Q1 – Q3) | 2.0 (0.0 – 6.0) | 4.5 (2.0 – 8.3) | 7.0 (5.0 – 9.3) | 1.0 (0.0 – 3.0) | 0.0 (0.0 – 0.0) | 0.0 (0.0 – 3.0) | MDD > SZ > AD & yHC & oHC; AD > yHC |
| *mean loneliness score (SD)* | 3.4 (3.6) | 4.8 (3.5) | 6.9 (3.0) | 2.1 (2.6) | 0.5 (1.1) | 1.5 (2.1) |  |

All patient groups had mild to moderate symptoms, and most patients were treated with psychotropic medication. All patients showed higher SFS and LON scores compared to their age-matched HC group (*p’s*<0.001), except for AD patients, who had similar LON scores compared to the HC-older group (*p*>0.05). Additionally, MDD patients had higher SFS and LON scores compared to SZ and AD patients (*p’s*<0.05).

The PRISM2 sample was highly comparable to its predecessor in the PRISM1 study, for instance in terms of age, years of education, disease severity, psychotropic medication use, and social functioning (8, 12). A few differences existed. First, the total sample size of PRISM2 was slightly larger compared to PRISM1 (8, 12), but due to the inclusion of an extra patient group (MDD), the sample size per patient group was smaller. Second, although the PRISM1 study had a higher percentage of male participants than PRISM2 (62.7% (8) versus 44.7%), within the AD and SZ patient groups and HC, the percentage of males was similar between studies. Finally, the mean LON score was higher in the resting-state PRISM2 sample compared to PRISM1 (8), again due to the inclusion of the MDD group. Characteristics of the diagnostic groups in the PRISM1 (8, 12) and PRISM2 studies were also highly similar (see Supplement).

### Social Dysfunction and DMN functional connectivity

The DMN maps found within the PRISM1 and PRISM2 sample (with and without MDD) are shown in Figure 4. A strong correlation (r=0.79) in DMN spatial structure was observed between the PRISM1 and PRISM2 sample (excluding MDD), and between the PRISM2 sample with and without MDD patients included (r=0.88), indicating a high degree of similarity in spatial organization of the DMN across cohorts.

**Figure 4.**
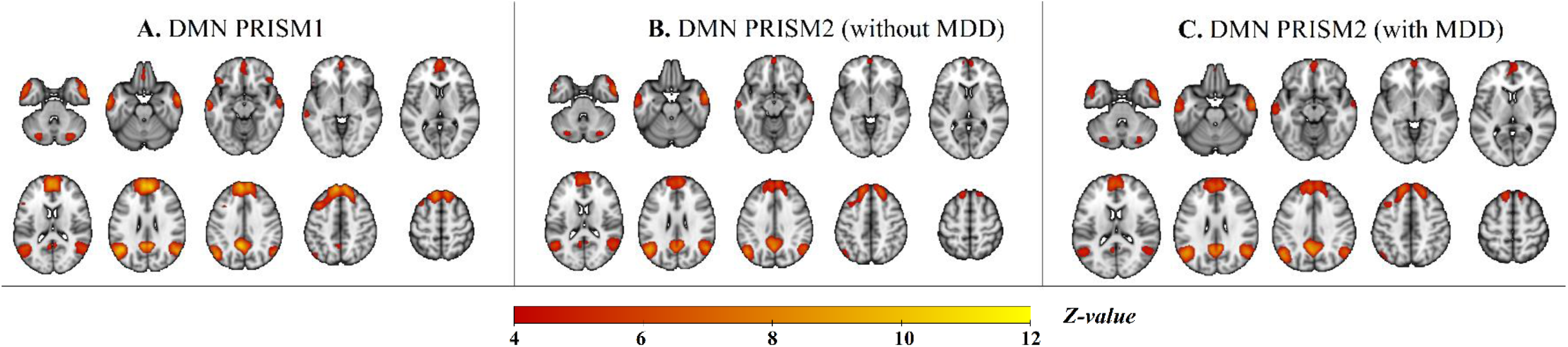
Axial view of the Default Mode Network constellation across the PRISM1 and PRISM2 studies. A): Default Mode Network (DMN) map across the PRISM1 sample. B): DMN map across the PRISM2 sample without (B) and with (C) major depressive disorder (MDD) patients included. The DMN maps comprised the posterior cingulate cortex, precuneus, medial prefrontal cortex, superior frontal gyrus, posterior inferior parietal lobe, temporoparietal junction, lateral temporal cortex, temporal pole, and cerebellum. The yellow-red scalar bar shows the connectivity strength (*Z*-value) within the DMN. The images are displayed in neurological convention.

Higher mean SFS+LON scores were associated with diminished DMN functional connectivity within the dmPFC, posterior cingulate cortex (PCC) and precuneus across SZ/AD/HC participants (*p*=0.009; x_max_=48, y_max_=43, z_max_=56; adjusted R^2^=0.25). In the total sample with MDD included, a similar association was found only within the dmPFC across participants (*p*=0.029; x_max_=51, y_max_=86, z_max_=58; adjusted R^2^=0.13) (Figure 5). After excluding the outlier (MDD patient), this association remained significant when focusing specifically on the effect site from the total sample (dmPFC) (*p*=0.003), but not in the whole-DMN analysis (*p*=0.12).

**Figure 5.**
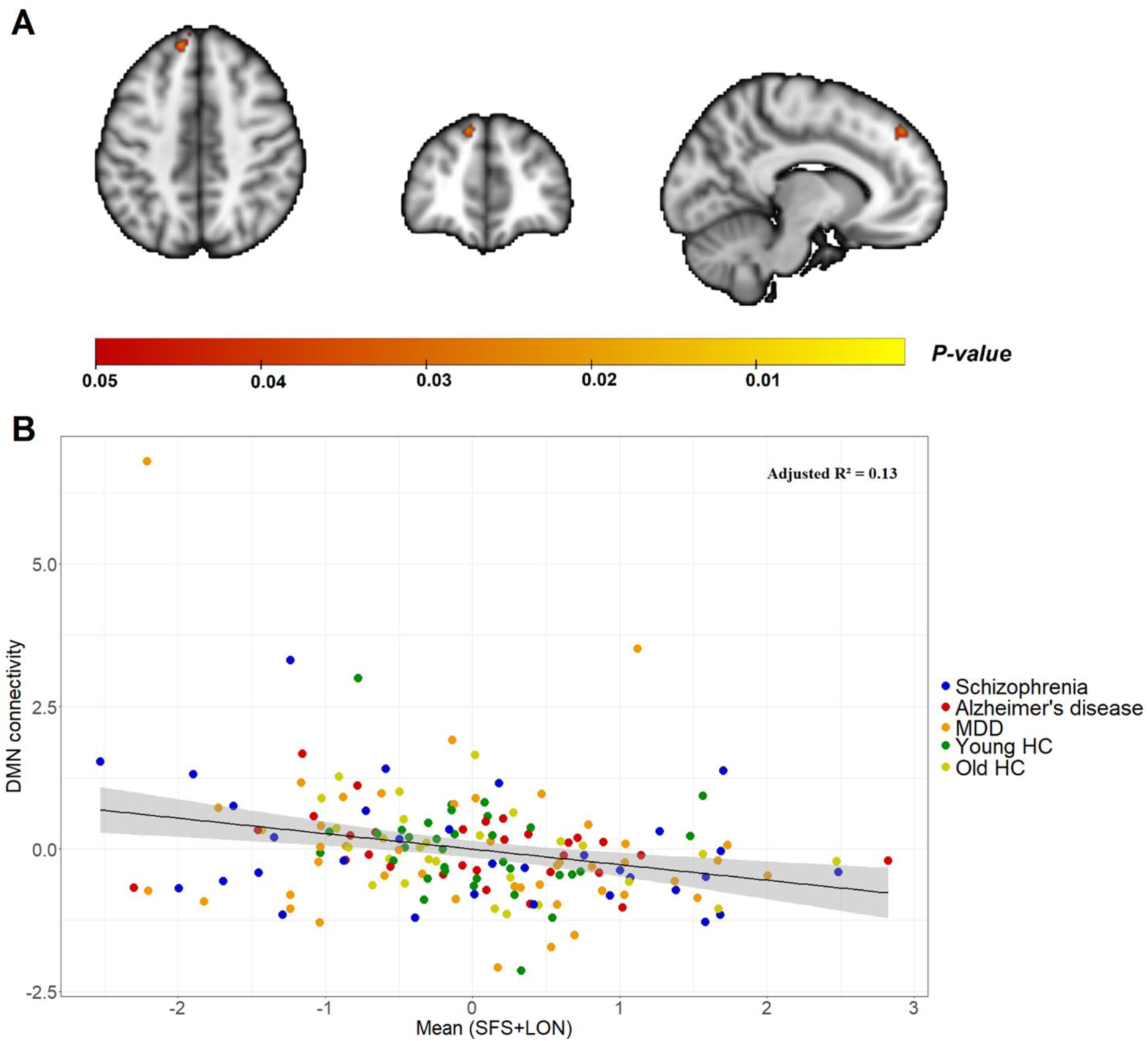
Social dysfunction and functional connectivity of the Default Mode Network. A): Significant cluster within the dorsomedial prefrontal cortex, wherein higher mean SFS+LON scores were associated with diminished functional connectivity across the sample (*p*=0.029; x_max_=51, y_max_=86, z_max_=58). The image is displayed in neurological convention. The yellow-red scalar bar shows the significance level (*P*-value) of this association within the significant cluster. B): The scatter plot visualizes this effect, wherein the mean connectivity within the significant cluster (y-axis) is plotted against the mean SFS+LON score (x-axis). The values on the y and x-axis are z-score residuals after correcting for clinical and sociodemographic factors. Higher positive values on the x-axis indicate more severe social dysfunction. The black solid line depicts the slope of the association, with the grey band indicating the 95% confidence interval of the slope. DMN = Default Mode Network, SFS = Social Functioning Scale, LON = de Jong-Gierveld Loneliness scale. MDD = Major depressive disorder. HC = Healthy controls.

Higher LON scores were associated with diminished DMN functional connectivity within the PCC and precuneus across SZ/AD/HC participants (*p*=0.014; x_max_=48, y_max_=43, z_max_=56; adjusted R^2^=0.24), but this association was not found in the analyses including the MDD patients (*p*>0.05). Unexpectedly, no associations were found between SFS scores and DMN functional connectivity across the sample with and without MDD participants (*p*’s>0.05). Additionally, no diagnosis x SFS and/or LON score interaction effects were observed for the findings (*p*’s>0.05) and patients (SZ/MDD/AD) showed higher DMN functional connectivity than HC (HC-younger & HC-older) (*p*<0.001) (Figure S1). Of note, no associations were observed between the SFS and/or LON scores and functional connectivity within the salience network or CEN across the sample with and without MDD participants (*p*’s>0.05). Finally, the mega-analysis, including PRISM1 and PRISM2 samples (N=317), revealed that higher mean SFS+LON scores were associated with diminished functional connectivity within large sections of both the dmPFC and rmPFC across SZ/MDD/AD/HC participants (*p<*0.001; x_max_=45, y_max_=87, z_max_=40; adjusted R^2^=0.47) (Figure 6). Similar associations were observed for the unique effect of LON scores (*p*=0.014; x_max_=45, y_max_=88, z_max_=40; adjusted R^2^=0.41), while diminished connectivity in the rmPFC was associated with higher SFS scores (*p*=0.004; x_max_=45, y_max_=90, z_max_=44; adjusted R^2^=0.35). The standardized beta estimates for most associations in the PRISM1 (8), PRISM2, and mega-analysis samples were comparable, particularly for mean SFS+LON and the unique effect of SFS scores, as shown in Figure 7.

**Figure 6.**
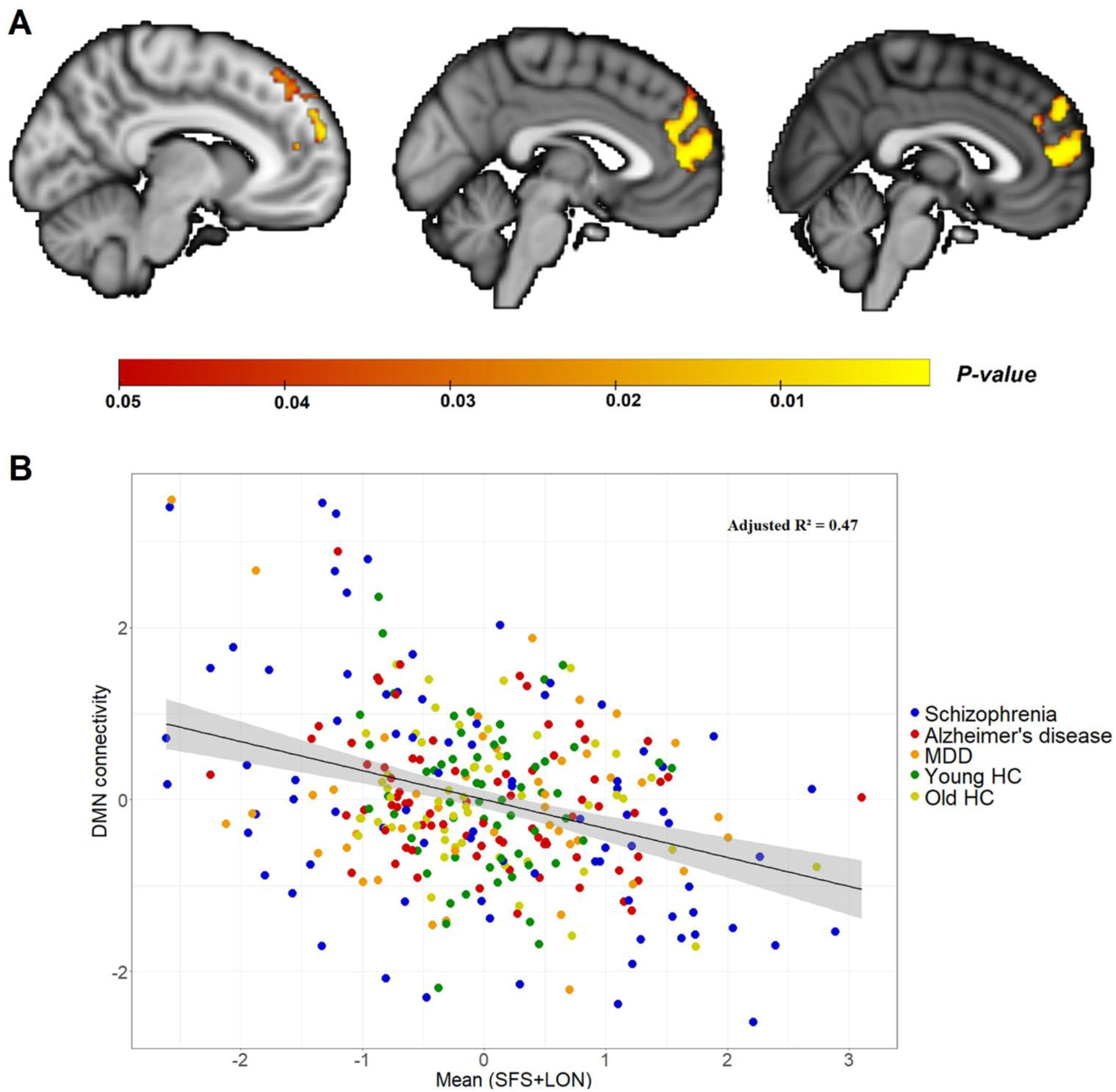
Social dysfunction and functional connectivity of the Default Mode Network in the mega-analysis. A): Significant cluster within the dorsomedial and rostromedial prefrontal cortex, wherein higher mean SFS+LON scores were associated with diminished functional connectivity across the sample (*p*<0.001; x_max_=45, y_max_=87, z_max_=40) (N=317). The image is displayed in neurological convention. The yellow-red scalar bar shows the significance level (*P*-value) of this association within the significant cluster. B): The scatter plot visualizes this effect, wherein the mean connectivity within the significant cluster (y-axis) is plotted against the mean SFS+LON score (x-axis). The values on the y and x-axis are z-score residuals after correcting for clinical and sociodemographic factors, and study sample. Higher positive values on the x-axis indicate more severe social dysfunction. The black solid line depicts the slope of the association, with the grey band indicating the 95% confidence interval of the slope. DMN = Default Mode Network, SFS = Social Functioning Scale, LON = de Jong-Gierveld Loneliness scale. MDD = Major depressive disorder. HC = Healthy controls.

**Figure 7.**
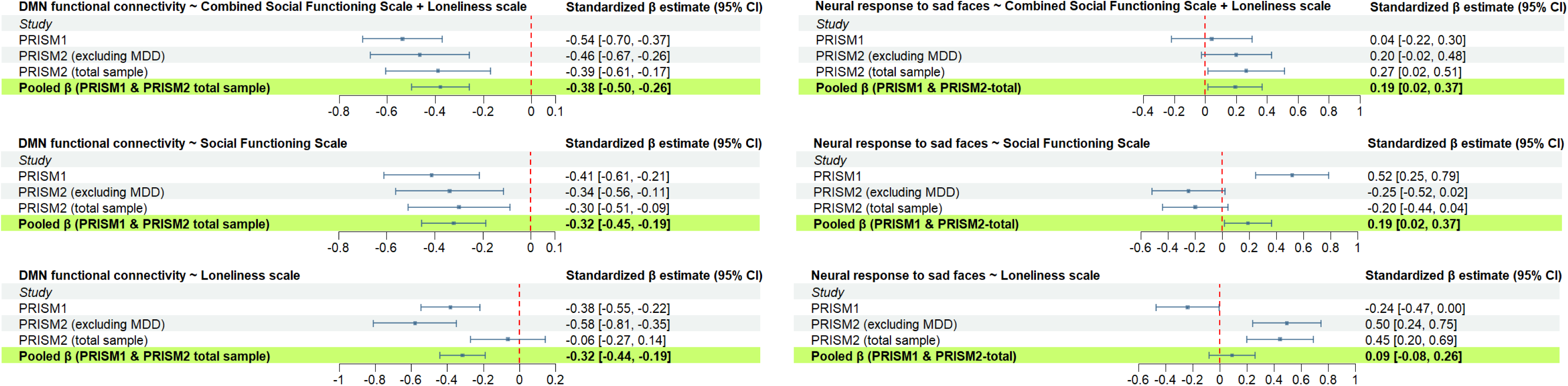
Forest plot of the social functioning indicators on Default Mode Network functional connectivity and activation. The forest plots on resting-state fMRI data are shown in the left panels, while the right panels show the forest plots of the implicit facial emotional processing (iFEP) fMRI task. On the top row the effect sizes (standardized beta estimates) of the mean Social Functioning Scale (SFS) + Loneliness scale (LON) effect are shown, while the effect sizes of the unique effect of the SFS and LON regressors are shown on the middle and on the bottom row, respectively. Effect sizes were calculated within the significant cluster found within each analysis. For resting-state, in case no significant cluster was found, effect sizes of associations between the unique effect of the SFS or LON regressor and Default Mode Network (DMN) connectivity in PRISM2 were calculated using the significant clusters found in the previous PRISM1 study (8). For the iFEP task, effect sizes for associations between mean SFS+LON and functional activation in response to sad faces in the PRISM1 study (12) and mega-analysis were calculated within the significant cluster found in the PRISM2 total sample including major depressive disorder (MDD). Effect sizes of the unique effect of the SFS or LON regressors were calculated within the significant cluster found in PRISM1 (within the DMN) (12) and PRISM2, respectively. The red vertical line represents the reference line equal to zero. The horizontal lines represent the 95% confidence intervals of the standardized beta estimates.

### Social dysfunction and DMN functional activation

Higher mean SFS+LON scores were associated with greater neural activation in a collection of DMN regions in response to sad faces across SZ/AD/HC participants (*p*=0.016; x_max_=47, y_max_=32, z_max_=45; adjusted R^2^=0.06). This included the dmPFC, PCC, precuneus, angular gyrus, middle temporal gyrus, and the lateral occipital cortex, and extended to larger sections of the frontal, parietal, temporal, and occipital cortices in the whole-brain analysis (*p*=0.019). In the total sample including MDD participants, a similar observation was found within the PCC and precuneus (*p*=0.018; x_max_=42, y_max_=34, z_max_=45; adjusted R^2^=0.05) (Figure 8), and extended to a larger section of these brain regions in the whole-brain analysis (*p*=0.028).

**Figure 8.**
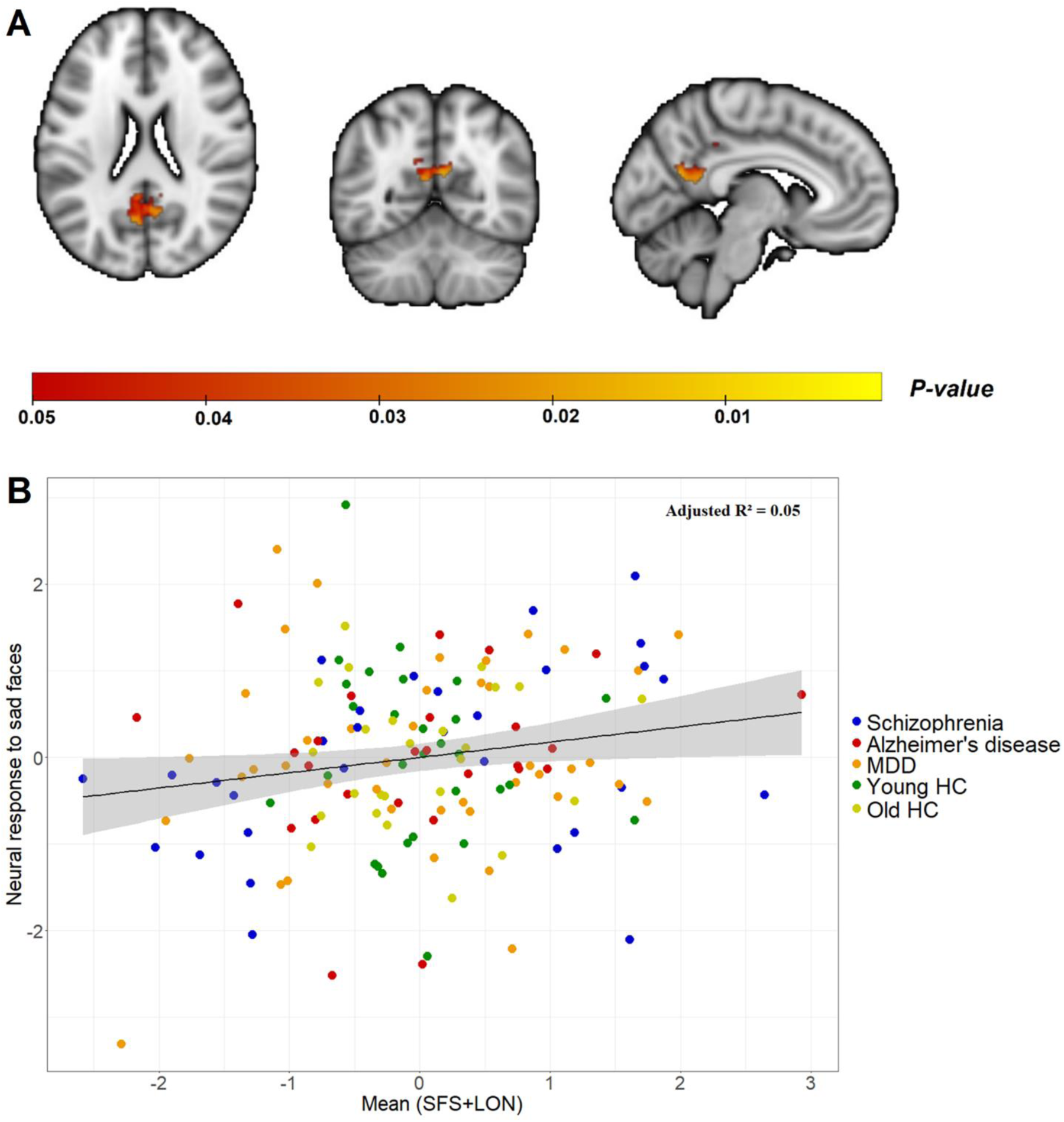
Social dysfunction and functional activation of the Default Mode Network in response to sad faces. A): Significant cluster within the posterior cingulate cortex and precuneus, wherein higher mean SFS+LON scores were associated with greater neural activation in response to sad faces across the sample (*p*=0.018; x_max_=42, y_max_=34, z_max_=45). The image is displayed in neurological convention. The yellow-red scalar bar shows the significance level (*P*-value) of this association within the significant cluster. B): The scatter plot visualizes this effect, wherein the functional activation within the significant cluster in response to sad faces (y-axis) is plotted against the mean SFS+LON score (x-axis). The values on the y and x-axis are z-score residuals after correcting for clinical and sociodemographic factors. Higher positive values on the x-axis indicate more severe social dysfunction. The black solid line depicts the slope of the association, with the grey band indicating the 95% confidence interval of the slope. SFS = Social Functioning Scale, LON = de Jong-Gierveld Loneliness scale. MDD = Major depressive disorder. HC = Healthy controls.

Unexpectedly, higher LON, but not SFS, scores were associated with greater neural activation in a largely overlapping set of brain regions - dmPFC, PCC, precuneus, angular gyrus, middle temporal gyrus, and the lateral occipital cortex - in response to sad faces across SZ/A D/HC participants (*p*=0.013; x_max_=47, y_max_=32, z_max_=45; adjusted R^2^=0.16), and to the PCC, precuneus and lateral occipital cortex across the sample with MDD participants included (*p*=0.015; x_max_=48, y_max_=32, z_max_=45; adjusted R^2^=0.09) (Figure S2). Post-hoc whole-brain analyses extended these associations to large sections of the frontal, parietal, temporal, and occipital cortices, and cerebellum, across SZ/AD/HC participants (*p*=0.025), and to the superior parietal lobe and lingual gyrus across SZ/MDD/AD/HC participants (*p*=0.023). No relationships were observed between higher SFS and/or LON scores and reduced brain activation in response to happy faces across the sample with and without MDD participants (*p’s*>0.05). The above findings were independent of diagnosis and no group differences in DMN responses to emotional faces were observed (*p’s*>0.05). Finally, the mega-analysis on combined PRISM1 and PRISM2 samples (N=291) revealed no associations between SFS and/or LON scores and neural processing of sad and happy faces within the DMN across SZ/MDD/AD/HC participants (*p’s>*0.05). However, the whole-brain analysis revealed higher SFS scores to be associated with reduced activation in the posterior insula, dorsolateral PFC, temporoparietal junction, precuneus, inferior frontal gyrus, precentral gyrus, and calcarine cortex, in response to happy faces across the participants (*p*=0.045; x_max_=72, y_max_=64, z_max_=42; adjusted R^2^=0.08). The standardized beta estimates for associations with sad faces in the PRISM1 (12), PRISM2, and mega-analysis samples showed greater variability than those from the resting-state analyses (Figure 7). The standardized beta estimates for associations with neural responses to happy faces (whole-brain) are shown in Figure S3. For completeness, the accuracy rates and reaction times for the iFEP task are shown in Table S4.

## Discussion

In the current study, we investigated the reproducibility of relationships between social dysfunction and diminished DMN intrinsic functional connectivity, as well as altered DMN functional activation in response to sad and happy faces, across SZ/AD patients and HC, and whether these relationships would extend to MDD patients. We replicated and generalized previous findings (8), linking greater social dysfunction to diminished DMN intrinsic functional connectivity. However, we did not observe similar associations between social dysfunction and DMN activation in response to emotional face stimuli, as in prior work (12). Our findings pinpoint altered DMN functional integrity as a novel treatment target for social dysfunction in diverse neuropsychiatric disorders, irrespective of diagnostic classifications.

### Social Dysfunction and DMN functional connectivity

The association between social dysfunction and diminished functional connectivity of the dmPFC across SZ/AD/HC participants was replicated, extended to the PCC/precuneus, and observed in MDD patients. The mega-analysis additionally showed diminished functional connectivity in large sections of the dmPFC and rmPFC. These brain regions are part of different DMN subsystems that are thought to be strongly interconnected and work in tandem to facilitate complex social behavior (37, 38). The rmPFC and PCC/precuneus form the core of the DMN, which is predominantly responsible for socio-cognitive information processing and self-referential thinking (37–41). The dmPFC is part of the dmPFC subsystem and is mainly implicated in mentalising (e.g., Theory of Mind) (37–40). Deficits in these cognitive functions are linked to social dysfunction in neuropsychiatric disorders (42–44) and may be amenable to interventions, such as repetitive transcranial magnetic stimulation or psychotropic medication, including psychedelics (40, 45–47). Our findings align with previous efforts in other neuropsychiatric disorders (e.g., autism spectrum disorder and ADHD) (14, 15), further supporting the paradigm shift towards understanding social dysfunction as a transdiagnostic dimensional construct across multiple modalities. Various factors may underly diminished DMN functional connectivity, including genetic influences and white-matter integrity. For instance, genes such as DRD2, which are expressed in prefrontal DMN regions (e.g., medial PFC), are strongly linked to sociability, and are associated with DMN functional connectivity within the medial PFC and PCC (48–50). Additionally, demyelination of the white-matter tracts (e.g., forceps minor) that connect these prefrontal DMN regions reduces DMN connectivity and impairs social behavior in mice (51). Similarly, the microstructural integrity of the forceps minor has been implicated in social dysfunction in humans (52). Ultimately, to identify stratified patient clusters relevant for treating social dysfunction, a multimodal (structural, functional, and connectional) approach may be taken, such as multimodal normative modeling (53).

### Social dysfunction and DMN functional activation

Unexpectedly, previous associations (12) between only the behavioral aspects of social dysfunction and altered DMN activation in response to sad and happy faces were not replicated. However, both the combined social dysfunction measures and perceived loneliness were associated with increased DMN activation in response to sad faces, indicating that resting-state functional connectivity may be a more robust measure compared to the currently used iFEP fMRI task when exploring brain-behavior relationships in social dysfunction. Inconsistencies of findings across functional activation studies in psychiatry have been documented (54). A potential explanation is that the greater complexity of the iFEP fMRI task, including the design and the cognitive processes involved, may lead to reduced reproducibility compared to resting-state fMRI, although this remains to be investigated. Nevertheless, consistent with previous work (12, 13), we provide further evidence that individuals with greater social dysfunction disproportionately perceive sad faces as salient socio-affective stimuli. Previous research has suggested that iFEP fMRI tasks may leave participants with sufficient cognitive capacity for self-reflective processing, as these tasks only require determining the gender of depicted faces via button-presses (55). Consequently, hyperactivity of the DMN, particularly within the PCC/precuneus, in response to sad faces might potentially also reflect heightened self-reflective processing of these emotions in the more socially dysfunctional individuals. Future research should explore whether down-regulating the neural hyperactivity in response to negative emotions, for example through cognitive behavioral therapy (56), leads to improved social functioning. This research may benefit from assessing social dysfunction as a broad concept that includes both behavioral and affective aspects, given this approach yielded the most robust findings in both the iFEP task and resting-state experiments (8, 10, 14). Finally, previous research linked social dysfunction to hyperactivity of the insula, inferior frontal gyrus, and calcarine cortex in response to negative emotions across psychotic disorders (57). Similarly, our mega-analysis showed that behavioral aspects of social dysfunction were associated with reduced activation in these brain regions in response to happy faces, cautiously suggesting that individuals with social dysfunction may also have deficits in processing rewarding or positive social information.

### Strengths and limitations

Using a highly similar data collection and analysis protocol, we successfully replicated a key part of the previous PRISM1 findings and extended these results to MDD patients (8, 12). By demonstrating that DMN connectional integrity is robustly associated with social dysfunction, independent of diagnosis, and diverging from disorder-specific findings by showing directional distinctiveness (increased vs. decreased) (see Supplement), our study may provide new directions for developing personalized treatment strategies. Nevertheless, several limitations should be acknowledged. Firstly, while the sample of the PRISM1 (8, 12) and the current study were highly similar, there were a few differences. The total sample size of PRISM2 was slightly larger compared to PRISM1 (8, 12), but the sample size per patient group was smaller. However, this is unlikely to alter the findings, which focused on the association between social dysfunction and DMN functional integrity across the sample. Additionally, the percentage of male participants in the PRISM1 study (8, 12) was higher compared to the PRISM2 study. Some evidence suggests that DMN connectivity patterns linked to social dysfunction are more strongly expressed in males than females, although these patterns were still significant in both genders in previous research (11). Besides, to mitigate potential gender effects, we corrected for gender in all imaging analyses. Secondly, due to the cross-sectional design, the directionality of the reported associations between social dysfunction and DMN functional integrity cannot be determined, although preliminary evidence may suggest a potential causal role of the DMN (51, 58–60). Finally, social dysfunction was assessed using self-report questionnaires, simplifying a complex domain and potentially introducing self-evaluation bias (61). Smartphone technology may potentially offer a more objective measure of social dysfunction by passively assessing social exploration and communication (62).

## Conclusion

In sum, the current study pinpoints DMN functional integrity, particularly within the medial PFC, as a robust cross-disorder target for developing more effective treatment strategies rooted in social behavior.

## Acknowledgements

The PRISM2 project has received funding from the Innovative Medicines Initiative 2 Joint Undertaking under grant agreement No 101034377. This Joint Undertaking receives support from the European Union’s Horizon 2020 research and innovation programme and EFPIA. This publication reflects only the author’s views and neither the IMI 2JU nor EFPIA nor the European Commission are liable for any use that may be made of the information contained therein. The IMI 2JU had no further role in study design; in the collection, analysis and interpretation of data; in the writing of the report; and in the decision to submit the paper for publication.

## Disclosures

GB is a consultant for Nxera Pharma and Neurocrine Biosciences and is a grant panel member for the Medical Research Council in the UK. GRD is owner of P1vital LTD that provides biomarkers tasks for clinical trials to both academia and the pharmaceutical industry. AB is an employee of P1vital Products Ltd. All other authors declare that they have nothing to disclose.

## Data availability statement

Data supporting the study findings are available from www.prism2-project.eu upon reasonable request to Prof. Martien Kas, provided dataset access policies are complied with.

