## Supplement for "Social dysfunction relates to altered default mode network functional integrity across neuropsychiatric disorders: A replication and generalization study"

***Supplement Braak et al.,***

### Methods and Materials

#### *Additional in- and exclusion criteria*

Additional cross-disorder exclusion criteria were: a) diagnosis of any other current psychiatric, neurological or somatic disorder that may affect the patient's ability to complete the study assessments, b) alcohol or drug abuse/dependence within previous 3 years, c) medically non-compliant in the management of their disease as judged by the investigator, d) no reliable study partner that can provide study data, e) severity of the illness that precludes the patient's ability to complete the study assessments.

Furthermore, schizophrenia (SZ) and Alzheimer's disease (AD) patients were excluded if they had a Quick inventory of Depressive Symptomatology Self-Rated (QIDS-SR) score ≥ 16, or if they had severe Parkinsonism as a consequence of antipsychotic medication (as assessed with a score ≥ 4 on the Extrapyramidal Symptom Rating Scale) (1). Additionally, AD patients with a history of strokes, as indicated by medical history or imaging results, were excluded. Major depressive disorder (MDD) patients were excluded if they received or were waitlisted for third line treatment, such as transcranial magnetic stimulation or electroconvulsive therapy.

The scores on the Mini-Mental State Examination - second edition for the healthy control (HC) participants had to be comparable to normative data according to age and years of education. Exclusion criteria for the HC groups were: a) a history of psychiatric Axis-I diagnosis or neurological diagnosis associated with cognitive impairment, b) somatic disorder that may affect the participant's ability to complete the study assessments, c) score > 5 on the QIDS-SR; d) current or prior use of antidepressant or anxiolytic medication including benzodiazepines; e) prescribed medication in the last 6 weeks that may affect the central nervous system and the participant's ability to complete the study assessments.

Finally, all participants (patients and HC) had to be right-handed or ambidextrous, free of any MRI contraindications, able to speak, read and write in the language in which psychometric tests were provided, not be socially withdrawn due to external circumstances (e.g. lack of access to transport) or comorbid medical disorder (e.g. lack of mobility), own a smartphone that is compatible with BeHapp installation, and finally not have participated in a study with an experimental drug within 90 days (or 5 times the half-life of the drug) or in two of those studies within 5 months, or in the previous PRISM1 study (2, 3).

#### *MRI data acquisition*

T1-weighted anatomical MRI scans were acquired with the following parameters: repetition time TR = 8.1 ms; echo time TE= 3.7 ms; flip angle 8°; 170 sagittal slices; no slice gap; field of view (FoV) 258 x 258 mm; 1 mm isotropic voxels; duration 5.29 min for Philips scanner; TR=2300 ms; TE=2.98 ms; flip angle 9°; 176 sagittal slices; no slice gap; FoV 256 x 256 mm; 1 mm isotropic voxels; duration 5.30 min for Siemens scanner.

Subsequently, T2*-weighted gradient-echo echoplanar imaging scans were acquired for resting-state with the following parameters: 240 whole-brain volumes; repetition time TR=2000ms; TE=30ms; flip angle 80°; 46 transverse slices; no slice gap; FoV 240x240 mm for Philips scanner and 210 x 210 mm for Siemens scanner; in-plane voxel size 3.0 x 3.0mm; slice thickness 3 mm; duration 8 min. Finally, for the iFEP task the following parameters were used: 340 whole-brain volumes; TR=2000ms; TE=30ms; flip angle 80°; 46 transverse slices; no slice gap; FoV 240 x 240 mm for Philips scanner and 210 x 210 mm for Siemens scanner; in-plane voxel size 3.0 x 3.0 mm; slice thickness 3 mm; duration 11.36 min for Philips and 11.29 min for Siemens.

#### *Imaging paradigms*

During the implicit facial emotional processing (iFEP) fMRI task, 120 emotional faces (sad, happy, fear) were presented in rapid succession in 12 separate blocks with 10 trials per block. The order of blocks is fixed across participants and each block starts with a 30s fixation cross to prepare for facial stimuli. To ensure task engagement, subjects were asked to identify whether the face was male or female with a button press. Each face appears for 100ms with a 2.9s inter-stimulus interval (ISI), allowing participants to respond during the ISI, and the next trial starts immediately after the ISI fixation cross. The task ends with a 30s fixation cross, bringing the total task duration to ~ 12 minutes. Stimuli were presented and responses were recorded using software developed by P1vital Products Limited. Participants practiced the task with different stimuli before scanning.

***fMRI Data Preprocessing***

Preprocessing consisted of nonbrain-tissue removal, motion correction, grand mean-based intensity normalization of the entire 4-D data set by a single scaling factor, slice timing correction, spatial smoothing with a 5 mm full width at half maximum Gaussian kernel, and temporal highpass filtering at 0.01 Hz (Gaussian-weighted least-squares straight line fitting). Functional data were registered to T1-weighted anatomical images, and subsequently to the 2-mm MNI standard space image, using boundary-based registration with 12 degrees-of-freedom and integrated distortion correction. On top of this, independent component analysis (ICA) based automated removal of motion artefacts was used for (micro)motion-related artefact removal. Additionally, white matter and cerebrospinal fluid signal removal was implemented to further clean the imaging data of noise (4). This was performed by creating masks of the white matter and cerebrospinal fluid for each participant, which were eroded at 90% to prevent partial voluming effects with grey matter, using FSL’s FAST (FMRIB's Automated Segmentation Tool) (5). These masks were then used to extract the average time series for white matter and cerebrospinal fluid, and these time series were used as nuisance regressors in the subject-level general linear model (GLM) analysis. This approach effectively removes physiological noise in functional data (6, 7). The maximum allowed mean displacement due to excessive head motion was set at 3 mm translation or 3º rotation in any direction. Resting-state fMRI data were available for 170 participants, 3 (SZ=1 and AD=2) participants were excluded because of excessive motion and poor imaging quality, leaving 167 participants for final resting-state imaging analyses. Regarding the iFEP task, fMRI data was available for 167 participants, but 8 (SZ=1; AD=5; HC-younger =2) were excluded due to excessive motion and poor imaging quality and 7 (SZ=1; MDD=1; HC-younger=3; HC-older=2) due to inaccurate time recording of stimulus onset, resulting in a final sample of 152 participants.

#### *Resting-state functional connectivity analyses*

A group-level probabilistic independent component analysis (ICA) was performed using MELODIC within FSL to obtain 20 temporally and spatially independent components from the preprocessed resting-state data (8). Subsequently, these sets of spatial maps from the group-average ICA were used to generate subject-specific versions of the spatial maps, and associated timeseries, using dual regression (8). The Default Mode Network (DMN) component was identified based on its topological architecture, as described in several overview papers (9, 10). Associations between diminished DMN functional connectivity and higher Social Functioning Scale (SFS) and/or De Jong-Gierveld Loneliness (LON) scale total scores were analyzed using FSL's Randomise permutation-testing tool with N=5000 permutations with automatic outlier-deweighting. Group comparisons of functional connectivity of the DMN were additionally performed using the same GLM model. Post-hoc analyses checked whether diagnosis x SFS and/or LON interaction effects could be identified. As a sensitivity analysis, we further checked whether findings were specific to the DMN, by rerunning the dimensional brain-social dysfunction analyses for the salience network and central executive network (CEN). The salience network and a right- and left-lateralized CEN were part of the 20 independent components obtained with MELODIC. To control for multiple testing, these statistical analyses were additionally Bonferroni corrected for the number of networks tested ($P$ 0.05/3 = 0.017).

#### *Implicit facial emotional processing task analyses*

Subject-level statistical analyses were performed using FMRIB’s Improved Linear Model (FILM) with local autocorrelation correction in FSL/FEAT (11). The regressors for each emotion category within the GLM were convolved with a double-gamma hemodynamic response function. To measure emotion-specific neural responses, two contrasts were tested in these subject-level GLM's: Sad vs. Fear & Happy (1 -0.5 -0.5) and Happy vs. Fear & Sad (1 -0.5 -0.5). Specific neural responses to fearful faces were not analyzed, since no associations between SFS and/or LON scores and neural responses to fearful faces were observed in the PRISM1 study (3). Using the subject-level statistical maps, group-level GLM analyses were performed using FSL's Randomise permutation-testing tool with N=5000 permutations with automatic outlier-deweighting (12, 13). Post-hoc analyses additionally checked whether diagnostic status x SFS and/or LON interaction effects could be identified.

### Results

#### *Patient characteristics*

Characteristics of the groups (SZ, AD, HC) in the current study were highly comparable to those in the previous PRISM1 studies, for example in terms of age, gender, years of education, disease severity, psychotropic medication use, and social functioning (2, 3). A few differences existed. First, SZ patients in the iFEP sample, but not in the resting-state sample, of PRISM2 had slightly lower SFS social dysfunction scores compared to the PRISM1 sample ($p<0.05$) (3). Second, the age of the HC-younger group in PRISM2 was higher compared to the PRISM1 ($p<0.05$), due to age-matching with the MDD group in PRISM2 and the different age-range of the HC-younger group (18-45 years in PRISM1 and 18-55 years in PRISM2) (2, 3).

#### *Diagnostics and DMN functional connectivity and activation*

Although this study was not designed to investigate alterations in DMN functional integrity as a function of diagnosis, the current study also examined differences in DMN functional connectivity and activation between diagnostic groups as a sensitivity test, similar to the previous PRISM1 studies (2, 3). The analyses revealed higher functional connectivity within the DMN among patients (SZ, MDD and AD combined) compared to HC (HC-younger and HC-older combined), comprising large sections of the prefrontal cortex, posterior cingulate cortex, precuneus and left angular gyrus ($p<0.001$) (Figure S3 in Supplementary data). No other group differences of DMN intrinsic functional connectivity or activation in response to the emotional face stimuli were observed (*p's*>0.05). Thus, similar to the previous PRISM1 studies (2, 3), the disorder-specific findings (and their absence) clearly diverge from the reported social dysfunction effects by showing directional distinctiveness (increased vs. decreased), further indicating that the findings are distinct and independent of diagnosis.

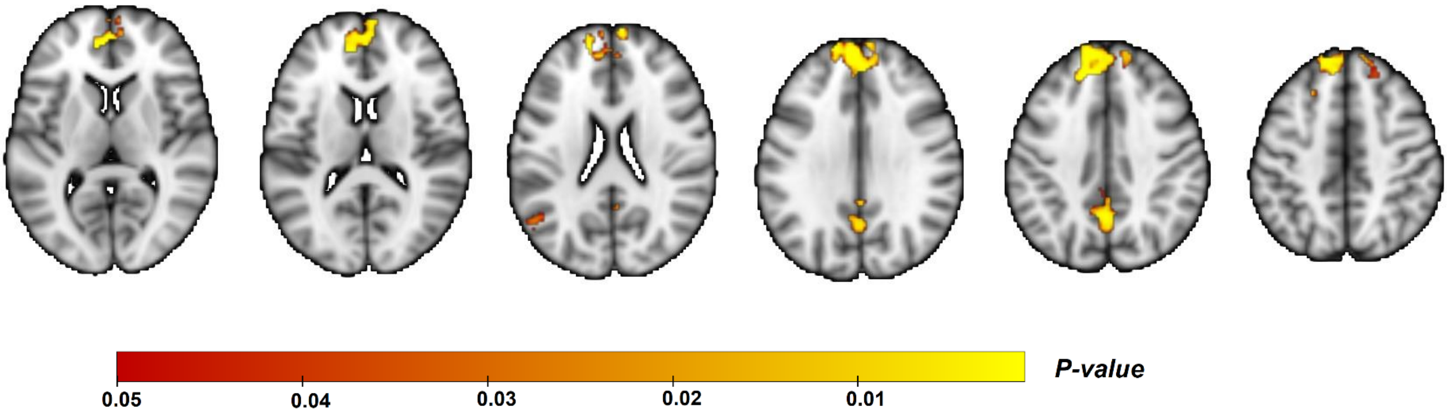


**Figure S1.** Diagnostics and functional connectivity of the Default Mode Network. Axial view of the significant cluster within the Default Mode Network (DMN), wherein patients (schizophrenia / Major Depressive Disorder / Alzheimer's disease) showed higher DMN functional connectivity compared to all healthy controls ($p<0.001$; $x_{max}=47$, $y_{max}=89$, $z_{max}=50$). The image is displayed in neurological convention. The yellow-red scalar bar shows the significance level (*P*-value) of this association within the effect site.

***Social dysfunction and functional activation***

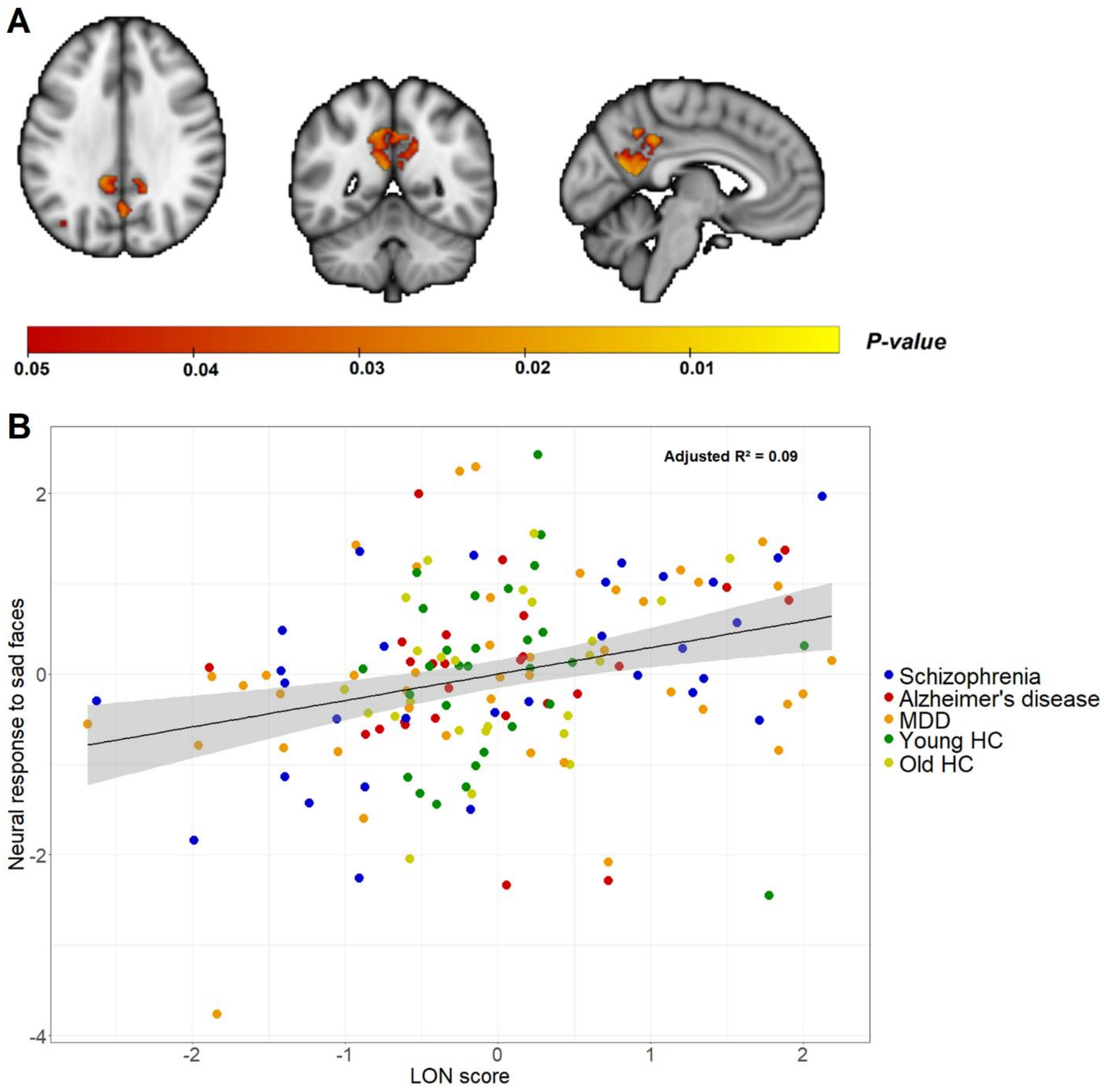


**Figure S2.** Perceived loneliness and functional activation of the Default Mode Network in response to sad emotional faces. A): Significant cluster within the posterior cingulate cortex, precuneus, and lateral occipital cortex, wherein higher perceived loneliness was associated with greater neural activation in response to sad faces across the sample ($p$=0.015; $x_{max}$=48, $y_{max}$=32, $z_{max}$=45). The image is displayed in neurological convention. The yellow-red scalar bar shows the significance level (*P*-value) of this association within the significant cluster. B): The scatter plot visualizes this effect, wherein the functional activation within the significant cluster in response to sad faces (y-axis) is plotted against the perceived loneliness score. The values on the y and x-axis are z-score residuals after correcting for clinical and sociodemographic factors. Higher positive values on the x-axis indicate more severe social dysfunction. The black solid line depicts the slope of the association, with the grey band indicating the 95% confidence interval of the slope. LON = de Jong-Gierveld Loneliness scale. MDD = Major depressive disorder. HC = Healthy controls.

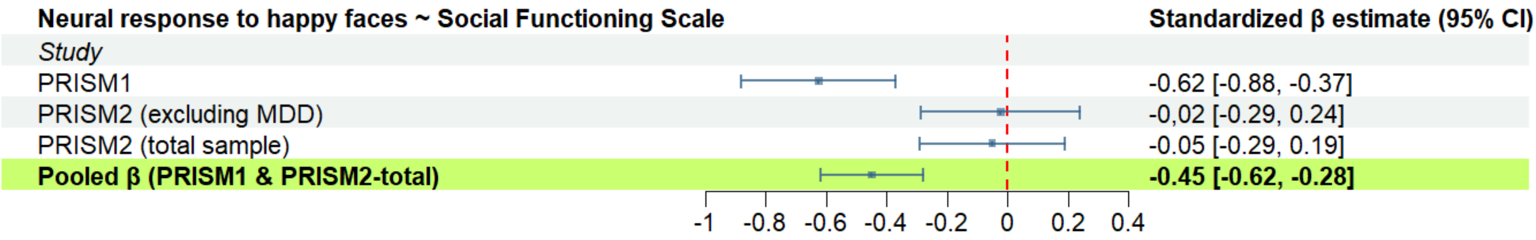


**Figure S3.** Forest plot of the social functioning indicators on functional activation in response to happy faces (whole-brain). The effect sizes (standardized beta estimates) of the association between higher Social Functioning Scale scores and reduced functional activation in response to happy faces are shown. Effect sizes were calculated within the significant cluster found within each analysis. Effect sizes in the PRISM2 samples were calculated using the same significant cluster found in the previous PRISM1 study (3). The red vertical line represents the reference line equal to zero. The horizontal lines represent the 95% confidence intervals of the standardized beta estimates. MDD = Major depressive disorder.

**Table S1. iFEP sample characteristics of each group**

| | Total Sample<br><br>N=152 | Schizophrenia (SZ)<br><br>N=30 | Major depressive disorder (MDD)<br><br>N=42 | Alzheimer's disease (AD)<br><br>N=26 | Young healthy controls (yHC)<br><br>N=29 | Old healthy controls (oHC)<br><br>N=25 | *Pairwise differences* |
|---|---|---|---|---|---|---|---|
| **Demographics** | | | | | | | |
| Age (years), median (Q1 – Q3) | 44.0 (31.8 – 66.0) | 32.5 (30.0 – 38.8) | 39.0 (31.0 – 48.8) | 72.0 (66.3 – 76.8) | 35.0 (28.0 – 44.0) | 70.0 (66.0 – 74.0) | AD & oHC > SZ & MDD & yHC |
| *mean age (SD)* | 47.7 (18.5) | 33.1 (7.0) | 38.3 (11.9) | 71.3 (5.8) | 36.8 (11.2) | 68.9 (5.3) | |
| Gender (% female) | 55.3% | 43.3% | 64.3% | 50.0% | 51.7% | 64.0% | $p > 0.05$ |
| Education (years), median (Q1 – Q3) | 15.0 (12.0 – 18.0) | 12.5 (10.0 – 16.0) | 15.0 (12.0 – 16.8) | 15.5 (12.0 – 19.5) | 17.0 (14.0 – 18.0) | 15.0 (13.0 – 18.0) | SZ < AD & yHC & oHC; MDD < yHC |
| *mean education (SD)* | 15.0 (3.8) | 12.9 (3.7) | 14.8 (3.5) | 15.3 (4.2) | 16.6 (3.2) | 15.5 (3.6) | |
| Number of individuals per scan site | | | | | | | |
| Amsterdam UMC | 50 | 10 | 13 | 11 | 7 | 9 | |
| Leiden UMC | 31 | 5 | 11 | 0 | 11 | 4 | |
| Spanish sites | 71 | 15 | 18 | 15 | 11 | 12 | |
| Specific disorder characteristics | | | | | | | |
| Psychotropic medication | | | | | | | |
| Antipsychotic (%) | 17.8% | 90.0% | 0.0% | 0.0% | 0.0% | 0.0% | |
| Antidepressant (%) | 23.7% | 10.0% | 64.3% | 23.1% | 0.0% | 0.0% | |
| Acetylcholinesterase inhibitor and/or NMDA receptor antagonist (%) | 9.9% | 0.0% | 0.0% | 57.7% | 0.0% | 0.0% | |
| Benzodiazepines (%) | 10.5% | 16.7% | 16.7% | 15.4% | 0.0% | 0.0% | |
| Antiepileptic (%) | 2.0% | 6.7% | 2.4% | 0.0% | 0.0% | 0.0% | |
| Other psychotropics (%) | 6.6% | 6.7% | 16.7% | 3.8% | 0.0% | 0.0% | |
| Severity disorder | | | | | | | |
| Positive symptoms, PANSS, mean (SD) | NA | 11.5 (2.5) | NA | NA | NA | NA | |
| Negative symptoms, PANSS, mean (SD) | NA | 14.9 (6.0) | NA | NA | NA | NA | |
| MMSE, median (Q1 – Q3) | NA | NA | NA | 25.0 (22.3 – 25.0) | 30.0 (29.0 - 30.0) | 30.0 (29.0 – 30.0) | AD > yHC & oHC |
| *mean MMSE (SD)* | | | | 23.6 (2.2) | 29.4 (0.7) | 29.3 (0.9) | |
| QIDS-SR, median (Q1 – Q3) | 3.0 (2.0 – 10.3) | 5.0 (3.0 – 11.0) | 14.0 (10.0 – 18.8) | 2.0 (1.3 – 4.5) | 2.0 (0.0 – 2.0) | 2.0 (2.0 – 3.0) | MDD > SZ > AD & yHC & oHC |

| | | | | | | | |
|---|---|---|---|---|---|---|---|
| *mean QIDS-SR (SD)* | 6.4 (6.2) | 6.6 (4.7) | 14.0 (5.2) | 3.1 (2.6) | 1.8 (1.7) | 2.0 (1.0) | |
| Social dysfunction scores | | | | | | | |
| Reversed SFS score, median (Q1 – Q3)*)* | 20.7 (14.7 – 30.1) | 23.5 (16.3 – 32.3) | 30.2 (25.7 – 36.3) | 20.8 (17.5 – 29.0) | 14.8 (9.2 – 17.3) | 13.6 (10.8 – 18.1) | MDD > AD & SZ > yHC & oHC |
| *mean SFS score (SD)* | 22.4 (10.7) | 24.7 (10.6) | 31.2 (8.7) | 23.1 (9.6) | 13.6 (5.0) | 14.6 (5.8) | |
| Loneliness score, median (Q1 – Q3) | 3.0 (0.0 – 6.0) | 4.5 (2.0 – 7.8) | 7.0 (5.0 – 9.8) | 1.0 (0.0 – 3.0) | 0.0 (0.0 – 1.0) | 0.0 (0.0 – 2.0) | MDD > SZ > AD & yHC & oHC |
| *mean loneliness score (SD)* | 3.5 (3.6) | 4.7 (3.5) | 7.1 (3.0) | 2.0 (2.5) | 0.6 (1.2) | 1.2 (1.6) | |

Mean and standard deviations (SD) are displayed for continuous variables. When assumptions were violated, median and Q1 – Q3 are also displayed. Chi-square tests were performed for categorical variables. For continuous variables, analysis of variance with Tukey´s method as a post-hoc test was performed. In case assumptions were violated, a Kruskal-Wallis test with Dunn´s test as a post-hoc test was performed. PANSS = Positive and Negative Syndrome Scale. MMSE = Mini-mental state examination. QIDS-SR = Quick Inventory of Depressive Symptomatology, Self-rated. SFS = Social Functioning Scale. A significance level of $p<0.05$ was considered statistically significant.

### Table S2. Resting-state mega-analysis sample characteristics

| | Total Sample N=317 | Schizophrenia (SZ) N=80 | Major depressive disorder (MDD) N=44 | Alzheimer's disease (AD) N=76 | Young healthy controls (yHC) N=62 | Old healthy controls (oHC) N=55 | *Pairwise differences* |
|---|---|---|---|---|---|---|---|
| **Demographics** | | | | | | | |
| Age (years), median (Q1 – Q3) | 44.0 (31.0 – 68.0) | 31.0 (26.8 – 37.0) | 39.0 (31.0 – 48.3) | 70.5 (65.0 – 75.0) | 30.5 (25.3 – 39.5) | 69.0 (63.0 – 73.0) | AD & oHC > SZ & MDD & yHC |
| *mean age (SD)* | 48.3 (19.4) | 31.8 (6.7) | 38.4 (11.7) | 69.7 (7.0) | 32.9 (10.2) | 67.9 (6.2) | |
| Gender (% female) | 47.0% | 33.8% | 61.4% | 46.1% | 48.4% | 54.5% | oHC & MDD > SZ |
| Education (years), median (Q1 – Q3) | 15.0 (12.0 – 18.0) | 14.0 (12.0 – 16.0) | 14.5 (12.0 – 16.3) | 15.0 (11.0 – 19.3) | 17.0 (15.0 – 18.0) | 16.0 (13.0 – 18.0) | SZ & MDD & AD < yHC: oHC > SZ |
| *mean education (SD)* | 15.2 (4.0) | 14.2 (3.9) | 14.5 (3.7) | 14.8 (4.5) | 16.8 (2.9) | 15.9 (4.3) | |
| Number of individuals per scan site | | | | | | | |
| UMC Utrecht | 20 | 12 | 0 | 0 | 3 | 5 | |
| Amsterdam UMC | 107 | 22 | 14 | 36 | 16 | 19 | |
| Leiden UMC | 48 | 10 | 11 | 2 | 14 | 11 | |
| Spanish sites | 142 | 36 | 19 | 38 | 29 | 20 | |
| Specific disorder characteristics | | | | | | | |
| Psychotropic medication | | | | | | | |
| Antipsychotic (%) | 23.3% | 90.0% | 0.0% | 2.6% | 0.0% | 0.0% | |
| Antidepressant (%) | 18.0% | 17.5% | 65.9% | 18.4% | 0.0% | 0.0% | |
| Acetylcholinesterase inhibitor and/or NMDA receptor antagonist (%) | 12.3% | 0.0% | 0.0% | 51.3% | 0.0% | 0.0% | |
| Benzodiazepines (%) | 8.8% | 13.8% | 15.9% | 10.5% | 0.0% | 3.6% | |
| Antiepileptic (%) | 2.8% | 8.8% | 2.3% | 1.3% | 0.0% | 0.0% | |
| Other psychotropics (%) | 4.4% | 6.3% | 15.9% | 1.3% | 0.0% | 1.8% | |
| Severity disorder | | | | | | | |
| Positive symptoms, PANSS, mean (SD) | NA | 11.2 (3.1) | NA | NA | NA | NA | |
| Negative symptoms, PANSS, mean (SD) | NA | 14.9 (5.9) | NA | NA | NA | NA | |
| MMSE, median (Q1 – Q3) | NA | NA | NA | 24.0 (23.0 – 25.0) | 30.0 (29.0 - 30.0) | 29.0 (28.0 – 30.0) | AD > yHC & oHC |
| *mean MMSE (SD)* | | | | 23.8 (2.0) | 29.4 (0.7) | 29.0 (1.1) | |

| | | | | | | | |
|---|---|---|---|---|---|---|---|
| QIDS-SR, median (Q1 – Q3) | 3.0 (2.0 – 8.0) | 7.5 (3.0 – 11.3) | 14.0 (10.0 – 19.0) | 3.0 (2.0 – 5.0) | 2.0 (1.0 – 3.0) | 2.0 (1.0 – 3.0) | MDD > SZ > AD > yHC & oHC |
| *mean QIDS-SR (SD)* | 5.5 (5.4) | 7.7 (5.1) | 13.9 (5.4) | 3.6 (2.6) | 2.0 (1.6) | 2.1 (1.2) | |
| Social dysfunction scores | | | | | | | |
| Reversed SFS score, median (Q1 – Q3)*)* | 20.6 (14.3 – 28.8) | 26.9 (21.4 – 34.1) | 30.2 (25.3 – 36.6) | 21.0 (16.9 – 28.0) | 14.2 (10.2 – 17.5) | 13.6 (9.9 – 16.8) | MDD & SZ > AD > yHC & oHC |
| *mean SFS score (SD)* | 21.9 (10.2) | 27.5 (9.6) | 31.2 (8.7) | 22.8 (8.0) | 14.1 (5.1) | 14.1 (6.3) | |
| Loneliness score, median (Q1 – Q3) | 1.0 (0.0 – 5.0) | 4.0 (1.0 – 8.0) | 7.0 (5.0 – 9.3) | 1.0 (0.0 – 3.0) | 0.0 (0.0 – 0.8) | 0.0 (0.0 – 2.5) | MDD > SZ > AD & yHC & oHC; AD > yHC |
| *mean loneliness score (SD)* | 3.0 (3.4) | 4.8 (3.7) | 6.9 (3.0) | 2.0 (2.3) | 0.5 (1.0) | 1.4 (1.9) | |

Mean and standard deviations (SD) are displayed for continuous variables. When assumptions were violated, median and Q1 – Q3 are also displayed. Chi-square tests were performed for categorical variables. For continuous variables, analysis of variance with Tukey´s method as a post-hoc test was performed. In case assumptions were violated, a Kruskal-Wallis test with Dunn´s test as a post-hoc test was performed. PANSS = Positive and Negative Syndrome Scale. MMSE = Mini-mental state examination. QIDS-SR = Quick Inventory of Depressive Symptomatology, Self-rated. SFS = Social Functioning Scale. A significance level of $p<0.05$ was considered statistically significant.

### Table S3. iFEP mega-analysis sample characteristics

| | Total Sample N=291 | Schizophrenia (SZ) N=76 | Major depressive disorder (MDD) N=42 | Alzheimer's disease (AD) N=66 | Young healthy controls (yHC) N=55 | Old healthy controls (oHC) N=52 | *Pairwise differences* |
|---|---|---|---|---|---|---|---|
| **Demographics** | | | | | | | |
| Age (years), median (Q1 – Q3) | 44.0 (30.5 – 67.0) | 31.0 (26.8 – 37.0) | 39.0 (31.0 – 48.8) | 71.0 (65.3 – 75.0) | 31.0 (26.0 – 40.0) | 69.0 (63.0 – 73.0) | AD & oHC > SZ & MDD & yHC |
| *mean age (SD)* | 48.0 (19.2) | 31.7 (6.7) | 38.3 (11.9) | 69.7 (6.7) | 33.3 (10.2) | 67.8 (6.3) | |
| Gender (% female) | 46.4% | 34.2% | 64.3% | 43.9% | 43.6% | 55.8% | oHC & MDD > SZ |
| Education (years), median (Q1 – Q3) | 15.0 (12.0 – 18.0) | 14.0 (11.0 – 16.0) | 15.0 (12.0 – 16.8) | 15.5 (11.0 – 20.0) | 17.0 (15.0 – 18.5) | 16.0 (13.0 – 18.0) | SZ < AD & yHC & oHC; MDD < yHC |
| *mean education (SD)* | 15.2 (4.1) | 14.0 (3.8) | 14.8 (3.5) | 14.8 (4.7) | 16.8 (2.9) | 16.1 (4.3) | |
| Number of individuals per scan site | | | | | | | |
| UMC Utrecht | 19 | 12 | 0 | 1 | 2 | 4 | |
| Amsterdam UMC | 98 | 21 | 13 | 29 | 15 | 20 | |
| Leiden UMC | 44 | 7 | 11 | 3 | 13 | 10 | |
| Spanish sites | 130 | 36 | 18 | 33 | 25 | 18 | |
| Specific disorder characteristics | | | | | | | |
| Psychotropic medication | | | | | | | |
| Antipsychotic (%) | 24.7% | 92.1% | 0.0% | 3.0% | 0.0% | 0.0% | |
| Antidepressant (%) | 17.9% | 15.8% | 64.3% | 19.7% | 0.0% | 0.0% | |
| Acetylcholinesterase inhibitor and/or NMDA receptor antagonist (%) | 10.7% | 0.0% | 0.0% | 47.0% | 0.0% | 0.0% | |
| Benzodiazepines (%) | 9.6% | 14.5% | 16.7% | 12.1% | 0.0% | 3.8% | |
| Antiepileptic (%) | 2.7% | 7.9% | 2.4% | 1.5% | 0.0% | 0.0% | |
| Other psychotropics (%) | 5.2% | 7.9% | 16.7% | 1.5% | 0.0% | 1.9% | |
| Severity disorder | | | | | | | |
| Positive symptoms, PANSS, mean (SD) | NA | 11.2 (3.2) | NA | NA | NA | NA | |
| Negative symptoms, PANSS, mean (SD) | NA | 14.8 (6.0) | NA | NA | NA | NA | |
| MMSE, median (Q1 – Q3) | NA | NA | NA | 24.0 (22.0 – 25.0) | 30.0 (29.0 - 30.0) | 29.0 (28.0 – 30.0) | AD > yHC & oHC |
| *mean MMSE (SD)* | | | | 23.7 (2.1) | 29.5 (0.7) | 29.1 (1.1) | |
| QIDS-SR, median (Q1 – Q3) | 3.0 (2.0 – 9.0) | 7.5 (3.0 – 11.3) | 14.0 (10.0 – 18.8) | 3.0 (2.0 – 5.8) | 2.0 (1.0 – 3.0) | 2.0 (1.0 – 3.0) | MDD > SZ > AD > yHC & oHC |
| *mean QIDS-SR (SD)* | 5.6 (5.4) | 7.7 (5.0) | 14.0 (5.2) | 3.8 (2.7) | 2.0 (1.6) | 2.0 (1.2) | |

| | | | | | | | |
|---|---|---|---|---|---|---|---|
| Social dysfunction scores | | | | | | | |
| Reversed SFS score, median (Q1 – Q3)) | 20.5 (14.4 – 29.4) | 28.0 (21.2 – 35.6) | 30.2 (25.7 – 36.3) | 21.0 (17.2 – 28.7) | 14.1 (10.3 – 17.4) | 13.3 (10.0 – 16.3) | MDD & SZ > AD > yHC & oHC |
| *mean SFS score (SD)* | 22.0 (10.3) | 27.9 (9.8) | 31.2 (8.7) | 22.9 (8.2) | 14.0 (5.0) | 13.5 (5.4) | |
| Loneliness score, median (Q1 – Q3) | 2.0 (0.0 – 5.0) | 4.5 (1.0 – 8.0) | 7.0 (5.0 – 9.8) | 1.5 (0.0 – 3.0) | 0.0 (0.0 – 1.0) | 0.0 (0.0 – 2.0) | MDD > SZ > AD & yHC & oHC; AD > yHC |
| *mean loneliness score (SD)* | 3.1 (3.4) | 4.9 (3.7) | 7.1 (3.0) | 2.1 (2.3) | 0.5 (1.1) | 1.3 (1.7) | |

Mean and standard deviations (SD) are displayed for continuous variables. When assumptions were violated, median and Q1 – Q3 are also displayed. Chi-square tests were performed for categorical variables. For continuous variables, analysis of variance with Tukey´s method as a post-hoc test was performed. In case assumptions were violated, a Kruskal-Wallis test with Dunn´s test as a post-hoc test was performed. PANSS = Positive and Negative Syndrome Scale. MMSE = Mini-mental state examination. QIDS-SR = Quick Inventory of Depressive Symptomatology, Self-rated. SFS = Social Functioning Scale. A significance level of $p<0.05$ was considered statistically significant.

### Accuracy and reaction times during the implicit facial emotional processing task

The accuracy rates and reaction times during the iFEP task for each group are shown in Table S4. HC-older participants had lower accuracy rates compared to SZ, MDD and HC-younger participants, but higher accuracy rates compared to AD patients (*p's*<0.05). The reaction times of HC-older participants were higher compared to AD patients and HC-younger participants (*p's*<0.05). Finally, the reaction times of MDD and SZ patients were higher compared to HC-younger participants (*p's*<0.05).

**Table S4. Accuracy and reaction times of the implicit facial emotional processing task**

| | Schizophrenia (SZ) N=30 | Major Depressive Disorder (MDD) N=42 | Alzheimer's disease (AD) N=26 | Young healthy controls (yHC) N=29 | Old healthy controls (oHC) N=25 | Pairwise differences |
|---|---|---|---|---|---|---|
| **Accuracy (%)** | 81.3% | 86.5% | 59.7% | 86.1% | 73.8% | MDD & yHC & SZ > oHC > AD |
| **Reaction time (ms)** | 868.1 | 830.6 | 791.3 | 751.9 | 899.1 | oHC > AD & yHC; MDD & SZ > yHC |

Mean accuracy (%) and reaction time (ms) of the implicit facial emotional processing task for each group. Kruskal-Wallis tests with Dunn´s test as post-hoc tests were performed.
